# Tropical butterflies lose central nervous system function in the cold from a spreading depolarization event

**DOI:** 10.1101/2023.05.31.543057

**Authors:** Mads Kuhlmann Andersen, Quentin Willot, Heath A. MacMillan

**Affiliations:** Department of Biology, Carleton University, Ottawa, ON, Canada K1S 5B6; Department of Biology, Aarhus University, Aarhus, Denmark 8000

**Keywords:** Spreading depression, cold tolerance, chill coma, Lepidoptera, CTmin

## Abstract

Insects are ectotherms and their physiological functions are therefore directly influenced by the environmental temperature. By extension, their ability to tolerate thermal extremes is directly linked to their thermal niche and distribution. Understanding the physiological mechanisms that limit insect thermal tolerance is therefore crucial for our ability to predict biogeography and range shifts. Recent studies on fruit flies and locusts suggest that the loss of coordinated movements at the critical thermal minimum is due to a loss of central nervous system function via a spreading depolarization. We hypothesized that a similar mechanism limits nervous function in other insect taxa. Here, we use electrophysiology to investigate whether the same spreading depolarization event occurs in the brain of butterflies exposed to stressful cold. Supporting our hypothesis, we find that exposure to stressful cold induced spreading depolarization in all species tested. This reinforces the idea that loss of central nervous function by a spreading depolarization is a common mechanism underlying the critical thermal minimum in insects. Furthermore, our results highlight how central nervous system performance is finely tuned to match species’ environments. Further research into the physiological mechanisms underlying the spreading depolarization event is likely to elucidate key mechanisms determining insect ecology.

## Introduction

Insects have limited ability to regulate their body temperature and environmental temperature is therefore one of the main factors influencing physiological functions and, by extension, their behaviour and fitness (1). Consequently, exposure to stressful thermal conditions constrain insect performance, and insects’ ability to tolerate thermal extremes is tightly associated with their fundamental niche range limits (2-4), a pattern that holds true for ectotherms in general (5, 6). This also applies to butterflies, where thermal tolerance traits have been used in attempts to model range limits and changes therein (7-10). Thus, gaining an integrative understanding of the physiological mechanisms limiting thermal tolerance and distribution of butterflies, is critical if we want to improve conservation efforts for some of the most charismatic, yet vulnerable species (11-13).

Several traits representing thermal tolerance limits can be measured in insects, the most common of which is the loss of movement at a temperature referred to as the critical thermal minimum or maximum (CT_min_ or CT_max_, see (14-16)). At low temperature, the gradual loss of function has been separated into multiple distinct, yet gradual, intertwined physiological events. Under the current consensus, CT_min_ refers to a gradual loss of coordinated movements quickly followed by paralysis as chill coma sets in (17-19). The physiological mechanism(s) underlying the CT_min_ and chill coma have been sought since cold-induced comas were first discovered in insects (20, 21). The ability to sustain movement is intrinsically linked to neuromuscular signalling, and research has therefore been focused on the temperature-sensitivity of neuronal and muscular physiology (22-25). Recent studies on fruit flies and locusts have found a strong correlation between the initial loss of coordination occurring at the CT_min_ temperature and a loss of central nervous system function caused by a spreading depolarization (17, 18, 26). Specifically, the spreading depolarization event silences central, integrating neurons due to a rapid surge in the brains extracellular K^+^ concentration (24, 27). The exact mechanisms underlying this event remain elusive, but multiple lines of evidence point towards impairment of ionoregulatory pumps and channels in perineurial glia that maintain the local ionic environment surrounding neurons (27-29).

Importantly, the proximate phenotype for spreading depolarization (i.e. CT_min/max_) is a wide-spread phenomenon in insects that is often used to capture discrepancies in thermal tolerance, predict species distributions, range expansions, and sensitivity to global climate change (2, 30, 31). However, so far, temperature-induced spreading depolarizations have only been demonstrated in fruit flies and locusts (17, 26); whether or not this neurophysiological limit is common to all insects is unclear. In the present study we sought to investigate whether the same spreading depolarization phenomenon occurs in another insect order by monitoring the transperineurial potential during a temperature ramp using a range of tropical butterfly species. Subsequently, we tested if this putative limit to organismal function (i.e. loss of central neural function) had any predictive power in terms of species biogeography.

## Materials and Methods

### Animal husbandry

All butterflies (see **Table 1**) were bought as pupae from a commercial supplier (LPS LLC Butterfly Import, Denver, CO, USA), housed under common garden conditions in flight cages at ∼ 30°C until emergence and left to mature in their flight cage or in a greenhouse (28-30°C) for 1-3 days before being used in experiments.

**Table 1.**
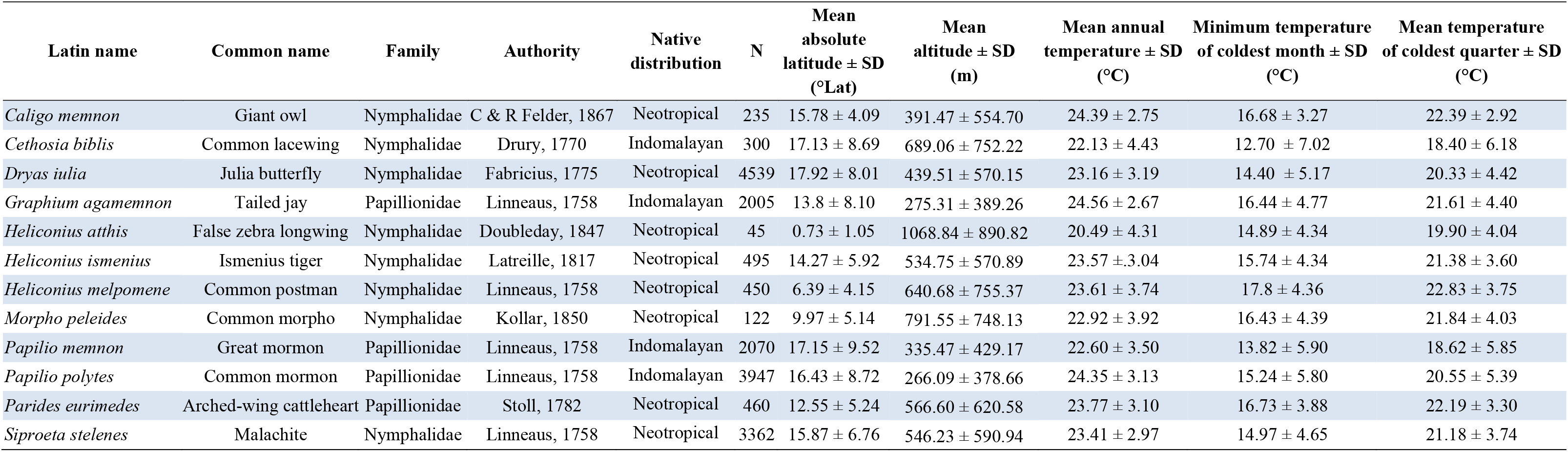
Description, classification, and biogeography of species used in this study.

### Experimental procedure

On the day of the experiment, individual butterflies were gently moved from a holding cage to the lab and immobilized in wax on top of a custom-built thermoelectrically cooled plate. Once immobilized, a small cut was made in the head immediately above the antennae, and a glass microelectrode was inserted into the brain through the blood-brain barrier with a micromanipulator (32). A reference electrode (Ag/AgCl) was inserted into the hemolymph through a small hole cut in the abdomen. The glass microelectrodes were fashioned from 1 mm borosilicate glass capillaries (1B100F-4, World Precision Instruments, Sarasota, FL, USA) pulled to a tip resistance of 5-7 MΩ in a Flaming-Brown P-1000 micropipette puller. Before use, glass electrodes were backfilled with 500 mmol L^-1^ KCl and placed in an electrode holder with an Ag/AgCl wire. Both glass and reference electrodes were connected to a Duo 773 two-channel intracellular/extracellular amplifier (World Precision Instruments, Sarasota, FL, USA), and the output was digitized with a PowerLab 4SP A/D converter (ADInstruments, Colorado Springs, CO, USA) and fed to a computer running Lab Chart 4 software (ADInstruments).

To approximate brain temperature, a type K thermocouple was placed immediately next to the head and at the same height as the tip of the glass microelectrode. Once ready, temperature was manually lowered from room temperature (∼22°C) by ∼ 1°C min^-1^ while the transperineurial potential (the electrical potential difference across the blood-brain-barrier) and head temperature was continuously recorded. Loss of CNS function caused by a spreading depolarization was identified when an abrupt negative shift in the transperineurial potential (∼20-45 mV in 5-10 s, see (32)) was observed, and the temperature of this event was estimated as the temperature at the half amplitude of the negative shift in potential (see **Fig. 1** for examples). After experiments, animals were dissected to determine their sex. Sample sizes are depicted in Fig. 2A.

**Figure 1.**
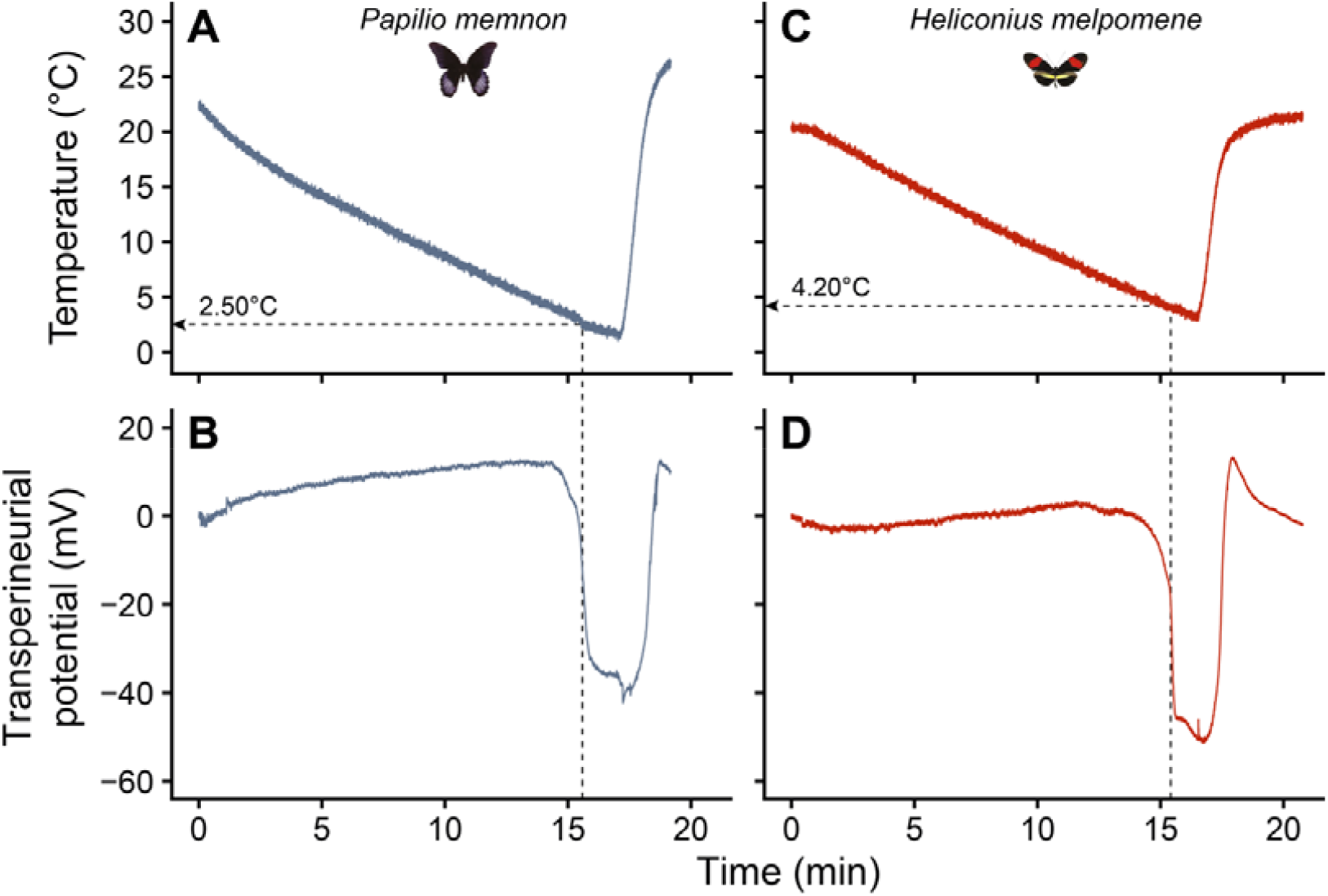
Example electrophysiological traces of the transperineurial potential during a cold ramp. To investigate if butterflies experience a spreading depolarization event during cooling, we monitored the transperineurial potential using glass microelectrodes inserted into the brain. As temperature is lowered (upper panels, A and C) there were only small changes in the transperineurial potential until at a certain temperature where it rapidly dropped > 25 mV in less than 10 s, indicative of a spreading depolarization having occurred (lower panels, B and D). Example traces are from *Papilio memnon* (A and B) and *Heliconius melpomene* (C and D), respectively.

**Figure 2.**
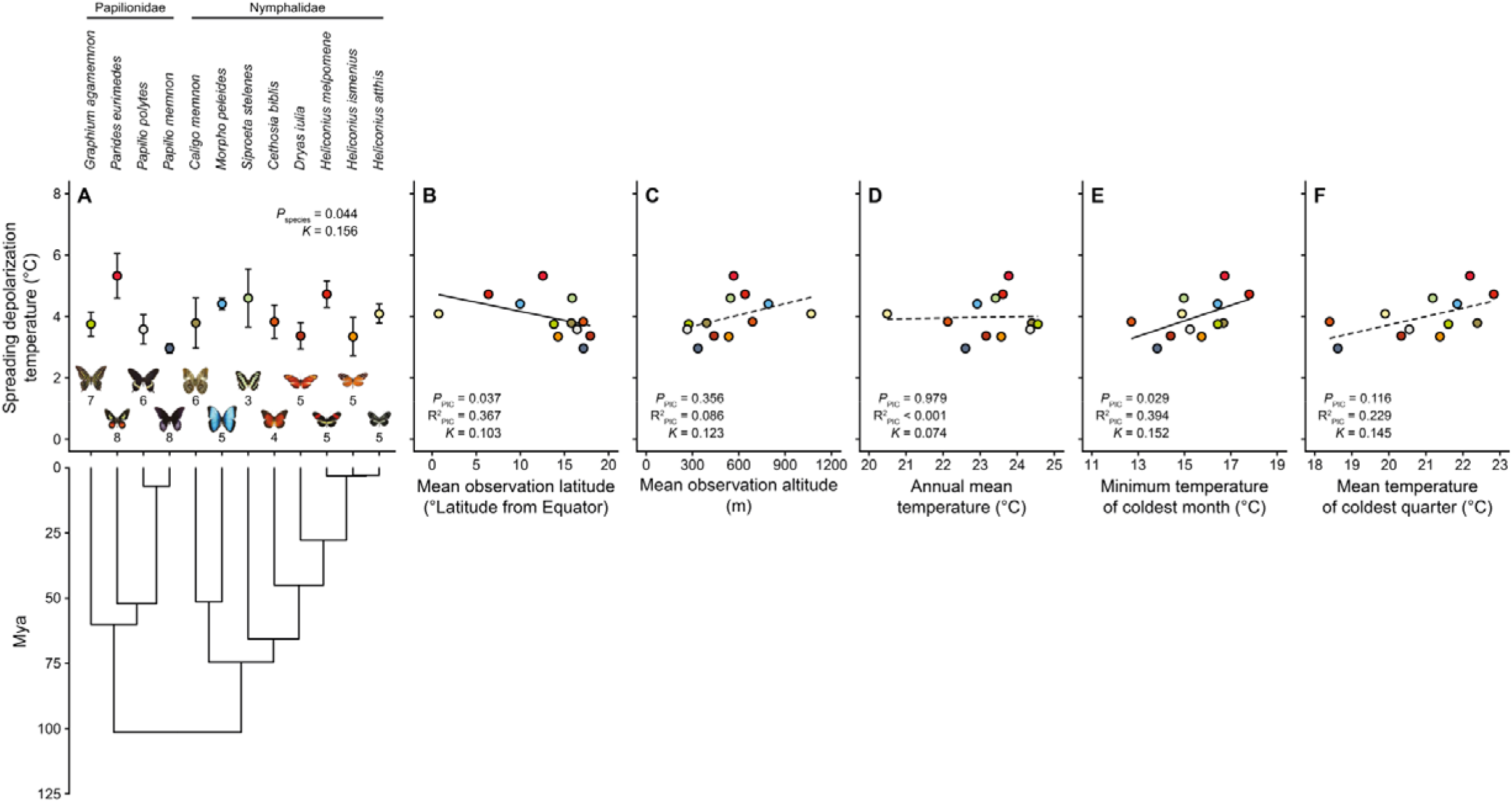
Temperatures leading to loss of central nervous function and correlations to biogeographical variables. (A) The temperature leading to a spreading depolarization-induced loss of CNS function, and its relation to species-specific (B) latitudinal distribution and the (C) mean altitude, (D) annual mean temperature, (E) minimum temperature of the coldest month, and (F) the mean temperature of the coldest quarter within their respective observed distributions. Sample sizes for spreading depolarization temperatures are depicted in panel A. Fully drawn linear regression lines indicate significant relationships while dashed lines indicate no relationship (based on PIC analyses). The phylogeny displayed under panel A shows species relatedness based on the phylogeny of Kawahara et al. (40).

### Biogeography and climate data

Information on the geographical distribution of all 12 butterflies species used in the present study was extracted from the Global Biodiversity Information Facility (GBIF, see Supplementary Table 1 for references to the species-specific datasets). However, we combined these observations with iNaturalist observations, as the GBIF had limited observations of two species (*Heliconius atthis* and *Caligo memnon*). Before obtaining climate data, the distribution datasets was cleaned up by 1) removing duplicate observations (two decimal accuracy in latitude and longitude) and 2) removing observations more than three standard deviations away from the mean latitude (33). This resulted in a total of 18030 observations (per species: N = 45-4539, average = 1502). After clean-up, distributional data was converted into spatial data points using the SpatialPoints() function from the “*sp*” package (34). Next, climate variables were downloaded from the WorldClim dataset (35) (http://worldclim.org/, September 30^th^ 2021) in raster format at a resolution of 2.5 min (∼ 20 km^2^) and imported into R using the getData() function from the “*raster*” package (36). Lastly, climate data was extracted from the spatial data points for each species using the extract() function (also from “*raster*”). Following extraction, mean and standard deviation of the climate variables was calculated for each species. For the purpose of investigating latitudinal distributions, we decided to use the absolute latitude to avoid distribution latitudes based on observations above and below the Equator (only for the sake of correlations; not for climate variable extractions) (6, 37). Given the focus here was cold tolerance, so we focused on the following biogeographical data: 1) mean latitude of observed distribution (degrees latitude from Equator), 2) mean altitude of the observed distribution, 3) annual mean temperature (in °C), 4) minimum temperature of the coldest month (in °C), 5) the mean temperature of the coldest quarter (in °C).

### Statistical analysis

All statistical analyses were performed using R software (v. 4.2.2, (38)). A two-way ANOVA was initially used to determine whether spreading depolarization temperature differed among species and sexes. No interaction was found between sex and species so it was removed from the analysis. An effect of sex was found, but sex was not included as a factor in subsequent analyses because this would result in extremely low sample sizes (see Supplementary Table 2) and because only females were sampled for one species (*Siproeta stelenes*).

To appropriately correlate spreading depolarization temperature to biogeographical data and climate variables, we wanted to assure that species relatedness wouldn’t confound our results (i.e. the close relationship between *Heliconius* species). Thus, we constructed a time-calibrated phylogeny with all 12 species in R using the “*ape*” package (39), based on the published nodes of divergence (40). Then, to correct our data for phylogeny, we calculated the phylogenetically independent contrasts (PIC) based on our constructed phylogeny using the pic() function from the “*ape*” package. Subsequently, PICs for spreading depolarization temperature were correlated to the biogeographical parameter PICs using linear regression through the origin (41). Phylogenetic signals (Bloomberg’s *K*) were calculated using the phylosig() function from the “*phytools”* package (42). The threshold for statistical significance was 0.05 for all analyses, and values references below refer to mean ± s.e.m. unless otherwise stated.

## Results and Discussion

### Cold-induced spreading depolarization in a subset of tropical butterflies

A cold-induced spreading depolarization was observed in all animals tested (64/ 64) in the same way it has been observed for locusts and fruit flies previously (32, 43); an abrupt negative shift in transperineurial potential of approximately 20-45 mV occurring over 5-10s which could be reversed when the animal was returned to a permissive temperature (example traces in **Fig. 1**). Spreading depolarization has also been observed in locusts and fruit flies when exposed to stressful heat and anoxia (43-45), and it has been experimentally induced in cockroaches (46). This phenomenon has been linked to the lower thermal limit for coordinated movements (i.e. the CT_min_, see (17, 18, 26)), and while we didn’t quantify behavioural CT_min_, we did note that butterflies ceased moving at temperatures around that leading to spreading depolarization. Thus, the loss of CNS function by a spreading depolarization has now been observed in four distantly related insect orders, encompassing both holo- and hemimetabolous groups. This generality suggests that spreading depolarization is a neurophysiological phenomenon shared for all insects, and by extension that the critical thermal minimum of all insects might be caused by spreading depolarization.

Spreading depolarization events are characterized by a collapse of ionoregulatory capacity resulting in a rapid surge in the extracellular K^+^ concentration within the central nervous system. The processes involved in ion balance regulation in the insect central nervous system are numerous, but a key transporter is the Na^+^/K^+^-ATPase (47), which was recently shown to be involved in modulating the temperature leading to loss of neural function (48, 49). Interestingly, some butterfly species (milkweed butterflies, Danainae) possess Na^+^/K^+^-ATPases with an unusual resistance to inhibitors (e.g. ouabain, see (50)) which are often used to induce spreading depolarization experimentally (28, 48). Similarly, butterflies generally possess an “unconventional” ion homeostasis consisting of a relatively high concentrations of K^+^ and low concentrations of Na^+^ in the hemolymph (51), of which the former is also a common way to induce spreading depolarization (46, 52). Future research on the curious ionoregulatory homeostasis of butterflies may provide key insights into to the mechanism underlying spreading depolarization, and their establishment as a new model of spreading depolarization represents a first step toward this goal.

### Correlations between cold-induced loss of neural function and biogeography

Temperatures leading to cold-induced spreading depolarization were species-specific (F_11,51_ = 2.0, *P* = 0.044) and ranged from 2.96 ± 0.15°C in *Papilio memnon* to 5.32 ± 0.73°C in *Parides eurimedes*, (**Fig. 2A**), however, the phylogenetic signal was low (*K* = 0.156). A low degree of interspecific variation in the lower thermal limit (chill coma onset temperature) of butterflies has been noted previously (53), however, this data was collected from wild-caught specimens of species with a wide geographical distribution. Behavioural chill coma temperatures estimated by Andersen et al. (51) revealed clear differences, however, the species compared were either domesticated (*Bombyx mori*), endemic to the neotropics (*Heliconius cydno*), or continent-spanning in distribution (*Manduca sexta*). We also found an effect of sex on the temperature leading to spreading depolarization (F_1,51_ = 7.8, *P* = 0.008) such that females generally experienced the event at lower temperatures than males. Sex-specific differences the critical thermal minimum have been noted previously (e.g. mosquitos (54)) but will not be discussed here.

To test whether temperatures leading to cold-induced spreading depolarization correlate with relevant biogeography we constructed a phylogeny of the species investigated to correct for relatedness. The phylogenetic signals were generally low (*K* < 0.2, see **Fig. 2**), but nevertheless accounted for in subsequent correlations. Doing so, we found significant correlations between the spreading depolarization temperature and 1) the mean distribution latitude (*P*_PIC_ = 0.037, R^2^_PIC_ = 0.367; **Fig. 2B**), and 2) the lowest temperature of the coldest month (*P*_PIC_ = 0.029, R^2^_PIC_ = 0.394; **Fig. 2E**). No relationship was found with the mean distribution altitude (**Fig. 2C**), the annual mean temperature (**Fig. 2D**), or the mean temperature of the coldest quarter (**Fig. 2F**). Strong relationships between cold tolerance measures and insect biogeography has been reported for several insect groups including, but not limited to, ants (55), fruit flies (2-4), and seed bugs (56), and meta-analyses have confirmed this trend on a global scales (5, 37). Nonetheless, this type of analysis does not account for microclimate variability and selection (57, 58), nor does it account for the ability of most adult lepidopterans to behaviourally thermoregulate (1), their dependence on appropriate host species (59, 60), ontogeny (61), or overwintering life stage (62). Furthermore, numerous butterfly species are known for their seasonal migration (63), adding further complexity to modelling seasonal range limits. Finally, it should also be noted that the temperatures leading to the cold-induced spreading depolarization are 11.5 ± 1.5°C (mean ± s.d., range = 9.1°C to 13.3°C) below the average lowest temperature of the coldest month, meaning that the butterflies are unlikely to experience it in nature. By extension cold-induced loss of central nervous function is unlikely to set the thermal limits to their poleward distributions, and instead they may be limited by other abiotic or biotic conditions that prevail outside of their native, tropical habitats (e.g. host availability). Nonetheless, low temperatures may still limit distribution *via* other mechanisms (e.g. degree-days (10)). Thus, to fully appreciate the factors determining butterfly distribution, and insect distribution in general, more complex models encompassing a wider range of species, species from varying climates, as well as a suite of abiotic and biotic factors are needed. Lastly, gaining an integrative understanding of mechanisms underlying other critical or subcritical limits to butterfly fitness and performance is likely to provide valuable predictive power in a world with a changing climate (11, 64).

## Acknowledgements

All animals used in the present study were kindly donated to us by Carleton University’s Annual Butterfly Show, and we would like to extend our utmost gratitude to Ed Bruggink who took care of them during their development. Butterfly images are used with permission from LPS LLC Butterfly Import, which we greatly appreciate.

## Author contributions

The study was conceived and designed by MKA and HAM. MKA performed the experiments, analyzed the data, and wrote the first draft. All authors contributed and made edits to the manuscript and approved the final version.

