## Supplementary material for "Tropical butterflies lose central nervous system function in the cold from a spreading depolarization event"

**Supplementary materials**

This document contains supplementary materials used in the study by Andersen et al. (*submitted*) titled ” Tropical butterflies lose central nervous system function in the cold from a spreading depolarization event”.

**Supplementary Table 1**

List of links to the GBIF distribution datasets used to extract biogeographical data for all 12 butterfly species.

| **Species** | **Date of download** | **Link** |
| --- | --- | --- |
| *Caligo memnon* | September 30, 2021 | <https://doi.org/10.15468/dl.ttfcm2> |
| *Dryas iulia* | September 30, 2021 | <https://doi.org/10.15468/dl.rrmwyg> |
| *Graphium agamemnon* | September 30, 2021 | <https://doi.org/10.15468/dl.k6yyb6> |
| *Heliconius atthis* | September 30, 2021 | <https://doi.org/10.15468/dl.wunp4f> |
| *Heliconius ismenius* | September 30, 2021 | <https://doi.org/10.15468/dl.fdhard> |
| *Heliconius melpomene* | September 30, 2021 | <https://doi.org/10.15468/dl.vwugdy> |
| *Morpho peleides* | September 30, 2021 | <https://doi.org/10.15468/dl.2wf85p> |
| *Papilio memnon* | September 30, 2021 | <https://doi.org/10.15468/dl.m74rz7> |
| *Papilio polytes* | September 30, 2021 | <https://doi.org/10.15468/dl.xnwhe9> |
| *Siproeta stelenes* | October 12, 2021 | <https://doi.org/10.15468/dl.mza7jx> |
| *Cethosia biblis* | October 12, 2021 | <https://doi.org/10.15468/dl.t7dnb9> |
| *Parides eurimedes* | October 14, 2021 | <https://doi.org/10.15468/dl.pv5k5c> |

**Supplementary Table 2**

This table contains all the measured spreading depolarization temperatures obtained for the 12 species used in this study.

| **Species** | **Sex**  **(M/F)** | **Spreading depolarization temperature**  **(°C)** |
| --- | --- | --- |
| *Caligo memnon* | F | 3.07 |
| *Caligo memnon* | M | 4.10 |
| *Caligo memnon* | F | 1.25 |
| *Caligo memnon* | F | 3.80 |
| *Caligo memnon* | M | 7.34 |
| *Caligo memnon* | M | 3.13 |
| *Cethosia biblis* | M | 4.48 |
| *Cethosia biblis* | M | 4.84 |
| *Cethosia biblis* | M | 3.56 |
| *Cethosia biblis* | F | 2.42 |
| *Dryas iulia* | M | 3.20 |
| *Dryas iulia* | M | 2.70 |
| *Dryas iulia* | F | 4.30 |
| *Dryas iulia* | F | 4.38 |
| *Dryas iulia* | M | 2.24 |
| *Graphium agamemnon* | M | 5.16 |
| *Graphium agamemnon* | F | 3.65 |
| *Graphium agamemnon* | F | 2.82 |
| *Graphium agamemnon* | M | 2.70 |
| *Graphium agamemnon* | F | 3.78 |
| *Graphium agamemnon* | M | 2.96 |
| *Graphium agamemnon* | M | 5.15 |
| *Heliconius atthis* | F | 4.65 |
| *Heliconius atthis* | F | 3.44 |
| *Heliconius atthis* | F | 3.23 |
| *Heliconius atthis* | M | 4.43 |
| *Heliconius atthis* | F | 4.68 |
| *Heliconius ismenius* | M | 5.64 |
| *Heliconius ismenius* | F | 2.50 |
| *Heliconius ismenius* | F | 2.05 |
| *Heliconius ismenius* | F | 3.02 |
| *Heliconius ismenius* | M | 3.49 |
| *Heliconius melpomene* | F | 3.72 |
| *Heliconius melpomene* | F | 4.25 |
| *Heliconius melpomene* | F | 5.40 |
| *Heliconius melpomene* | M | 4.20 |
| *Heliconius melpomene* | M | 6.03 |
| *Morpho peleides* | F | 3.94 |
| *Morpho peleides* | M | 3.98 |
| *Morpho peleides* | M | 4.69 |
| *Morpho peleides* | F | 4.65 |
| *Morpho peleides* | M | 4.78 |
| *Papilio memnon* | F | 2.81 |
| *Papilio memnon* | M | 2.97 |
| *Papilio memnon* | M | 2.50 |
| *Papilio memnon* | F | 2.30 |
| *Papilio memnon* | M | 3.18 |
| *Papilio memnon* | M | 3.54 |
| *Papilio memnon* | F | 2.87 |
| *Papilio memnon* | F | 3.49 |
| *Papilio polytes* | M | 3.37 |
| *Papilio polytes* | F | 3.46 |
| *Papilio polytes* | F | 1.78 |
| *Papilio polytes* | M | 3.13 |
| *Papilio polytes* | M | 5.07 |
| *Papilio polytes* | M | 4.63 |
| *Parides eurimedes* | M | 4.17 |
| *Parides eurimedes* | M | 6.80 |
| *Parides eurimedes* | F | 5.59 |
| *Parides eurimedes* | M | 6.86 |
| *Parides eurimedes* | F | 3.17 |
| *Siproeta stelenes* | F | 2.73 |
| *Siproeta stelenes* | F | 5.77 |
| *Siproeta stelenes* | F | 5.28 |
